# Pan-cancer analysis reveals embryonic and hematopoietic stem cell signatures to distinguish different cancer subtypes

**DOI:** 10.1101/2023.07.05.547742

**Authors:** Jiali Lei, Jiangti Luo, Qian Liu, Xiaosheng Wang

## Abstract

**Purpose:** Stem cells-like properties in cancer cells may confer cancer development and therapy resistance. With the advancement of multi-omics technology, the multi-omics-based exploration of cancer stemness has attracted certain interests. However, subtyping of cancer based on the combination of different types of stem cell signatures remains scarce.

**Methods:** In this study, 10,323 cancer specimens from 33 TCGA cancer types were clustered based on the enrichment scores of six stemness gene sets, representing two types of stem cell backgrounds: embryonic stem cells (ESCs) and hematopoietic stem cells (HSCs).

**Results:** We identified four subtypes of pan-cancer, termed StC1, StC2, StC3 and StC4, which displayed distinct molecular and clinical features, including stemness, genome integrity, intratumor heterogeneity, methylation levels, tumor microenvironment, tumor progression, chemotherapy and immunotherapy responses, and survival prognosis. This subtyping method for pan-cancer is reproducible at the protein level.

**Conclusion:** Our findings indicate that the ESC signature is an adverse prognostic factor, while the HSC signature and ratio of HSC/ESC signatures are positive prognostic factors in cancer. The ESC and HSC signatures-based subtyping of cancer may provide insights into cancer biology and clinical implications of cancer.

## Introduction

Stem cells, characterized by two critical abilities: self*-*renewal and differentiation, are the foundation for various organs and tissues [1]. There are different types of stem cells, including embryonic stem cells (ESCs), adult stem cells (ASCs), and induced pluripotent stem cells (iPSCs). ESCs are cells derived from very early embryos called blastocysts, which are pluripotent. In contrast, ASCs, also known as tissue-specific stem cells or somatic stem cells, generate specific cell types for specific tissues or organs and are not pluripotent [1]. ASCs mainly include hematopoietic stem cells (HSCs), mesenchymal stem cells, neural stem cells, epithelial stem cells, and skin stem cells. Among them, HSCs are the most representative ASCs that provide a standard model for studying tissue-specific stem cells [1]. HSCs are responsible for the generation of all types of blood cells, including red blood cells, white blood cells, and platelets. White blood cells, also known as leukocytes, are the cells associated with the immune system throughout the body, including lymphocytes, monocytes, neutrophils, eosinophils, basophils, and macrophages.

Cancer stem cells (CSCs) refer to cancer cells with stem cells-like characteristics. CSCs were initially discovered in hematopoietic malignancies [2] and later were found in various solid tumors, such as pancreatic cancer [3], melanoma [4], breast cancer [5], and head and neck cancer [6]. CSCs often constitute a small fraction of cancer cells, which are characterized by continuous self-renewal and proliferation, dedifferentiation, promoting cancer progression, and conferring therapy resistance [7].

CSCs are often endowed with dysregulated proliferative pathways or oncogenic mutations to maintain tumor growth [8]. Furthermore, different levels of stemness of cancer cells contribute to the intratumor heterogeneity (ITH) [9], and this heterogeneity allows drugs to kill a proportion of but not all cancer cells, a key factor responsible for therapy resistance and relapse in cancer. CSCs express many genes in common with early ESCs, such as *OCT4*, *NANOG*, and *SOX2* [10], most of which are transcriptional regulators of ESCs. These genes may synergistically regulate tumor cell self-renewal and proliferation [11]. Furthermore, because HSCs are the origin of all lymphocytes and myeloid cells [12], they play important roles in the immune regulation of various diseases, including cancer [13]. A previous study has suggested that hematopoietic malignancies and solid tumors may utilize the hematopoietic stem cell niche of the bone marrow to promote tumor growth and suppress antitumor immunity [14]. Another study has shown that hematopoietic stem and progenitor cells (HSPCs) are involved in tumor progression [15].

With the recent advancement of multi-omics technology, abundant cancer-associated genomics, epigenomics, transcriptomics, and proteomics data have emerged, such as The Cancer Genome Atlas (TCGA, https://cancergenome.nih.gov/) and The International Cancer Genome Consortium (ICGC, https://dcc.icgc.org/). These data have been widely utilized to explore tumor characteristics across various cancer types, such as tumor immunity [16], tumor stemness [17], tumor metabolism [18], and ITH [19]. For example, Thorsson et al. identified six immune-specific subtypes of the pan-cancer of 33 TCGA cancer types based on a transcriptomic analysis [20]. Another pan-cancer analysis of transcriptomes in 33 TCGA cancer types identified three metabolic expression subtypes with distinct clinical outcomes [18]. Hoadley et al. performed clustering of around 10,000 cancer samples from the 33 TCGA cancer types based on data of aneuploidy, DNA hypermethylation, mRNA, miRNA, and protein expression levels and identified 28 clusters significantly associated with histology, tissue type, or anatomic origin [21]. The multi-omics-based exploration of tumor stemness has attracted certain interests. For example, multi-omics analyses have shown that tumor stemness is associated with dedifferentiated oncogenic phenotype, immunosuppression, ITH, metastasis, and drug resistance [7, 17]. Transcriptomes-based stemness scores have been utilized to classify cancers, including bladder urothelial carcinoma (BLCA) [22], lung adenocarcinoma (LUAD) [9], and glioblastoma multiforme (GBM) [23]. Despite these previous studies, the omics-based investigation of different stem cell backgrounds in pan-cancer or individual cancer types remains lacking. Furthermore, the omics-based exploration of tumor stemness at the single-cell level remains inadequate, although there already been a wealth of data on single-cell omics available for a variety of cancers [24–28].

In this study, we performed a clustering analysis of 10,323 specimens representing the 33 TCGA cancer types based on transcriptomes-based scores of six stemness gene sets, which represent two types of stem cell backgrounds, namely ESCs and HSCs. This analysis identified four stemness subtypes of pan-cancer. We further comprehensively compared the molecular and clinical features among these subtypes, as well as their associations with the response to immunotherapy and chemotherapy.

## Materials and Methods

### Data acquisition

We downloaded data of gene expression profiles (RSEM normalized and batch effects adjusted), somatic mutation profiles (level 3) and clinical features for the TCGA Pan-Cancer (PANCAN) cohort consisting of 33 TCGA cancer types from UCSC Xena (https://xenabrowser.net/datapages/). The 33 cancer types included adrenocortical carcinoma (ACC), BLCA, breast invasive carcinoma (BRCA), cervical squamous cell carcinoma and endocervical adenocarcinoma (CESC), cholangiocarcinoma (CHOL), colon adenocarcinoma (COAD), lymphoid neoplasm diffuse large B-cell lymphoma (DLBC), esophageal carcinoma (ESCA), GBM, head and neck squamous cell carcinoma (HNSC), kidney chromophobe (KICH), kidney renal clear cell carcinoma (KIRC), kidney renal papillary cell carcinoma (KIRP), acute myeloid leukemia (LAML), brain lower grade glioma (LGG), liver hepatocellular carcinoma (LIHC), LUAD, lung squamous cell carcinoma (LUSC), mesothelioma (MESO), ovarian serous cystadenocarcinoma (OV), pancreatic adenocarcinoma (PAAD), pheochromocytoma and paraganglioma (PCPG), prostate adenocarcinoma (PRAD), rectum adenocarcinoma (READ), sarcoma (SARC), skin cutaneous melanoma (SKCM), stomach adenocarcinoma (STAD), testicular germ cell tumors (TGCT), thyroid carcinoma (THCA), thymoma (THYM), uterine corpus endometrial carcinoma (UCEC), uterine carcinosarcoma (UCS), and uveal melanoma (UVM). In addition, we downloaded 15 normalized protein expression profiles in 12 cancer types (TCGA-BRCA, TCGA-COAD, TCGA-OV, BRCA, COAD, OV, GBM, HNSC, LUAD, LUSC, PAAD, KIRC, UCEC, STAD, LIHC) from the Clinical Proteomic Tumor Analysis Consortium (CPTAC, https://proteomics.cancer.gov/programs/cptac/) and the International Cancer Proteogenome Consortium (ICPC, https://icpc.cancer.gov/portal/). We also downloaded three single-cell RNA sequencing (scRNA-seq) datasets, namely GSE75688 for BRCA [28], GSE131309 for SARC [29], and GSE89567 for GBM [24], from the NCBI gene expression omnibus (GEO) (https://www.ncbi.nlm.nih.gov/geo/). Besides, we downloaded a scRNA-seq dataset for renal cell carcinoma (RCC) from a recent publication [30]. We obtained eight immunotherapy-related datasets for eight cancer types (READ, non-small cell lung carcinoma (NSCLC), LIHC, ESCA, urothelial cancer (UC), triple-negative breast cancer (TNBC), STAD, SKCM) from GEO and recent publications [31–38]. The immunotherapy-related datasets involved gene expression profiles and clinical information on immune checkpoint blockade (ICB) treatment in cancer patients. A summary of these datasets is shown in Supplementary Table S1.

### Data processing and quality control

The gene expression values were preprocessed and normalized in the single cell transcriptomes, except the BRCA scRNA-seq dataset, in which they were raw transcripts per million (TPM). For the BRCA scRNA-seq dataset, we performed data preprocessing and quality control prior to subsequent analyses. First, we replaced the TPM values less than one with zero. Second, all TPM values (*x*) were transformed by log_2_(*x*+1). Third, we removed the genes with expression values of zero across all single cancer cells. For the other three scRNA-seq datasets, we also preprocessed them following the third step.

Proteomics data were processed by the publication [39] at the gene level rather than at the protein isoform level. The names of rows in the protein expression matrix refer to the protein-coding genes. For the pan-caner analysis, we merged 15 protein expression profiles into an expression matrix with the function “merge ()” in the R package “base.” We adjusted for batch effects and normalized combined data using the ‘‘normalizeBetweenArrays” function in the R package ‘‘limma.”

### Collection of stemness signatures

We collected six stemness signatures (or gene sets) from the StemChecker webserver (http://stemchecker.sysbiolab.eu). Among these stemness signatures, three are markers of ESCs, including Hs_ESC_Assou, Hs_ESC_Bhattacharya, and Hs_ESC_Wong; the other three are markers of HSCs, including Hs_HSC_Huang, Hs_HSC_Novershtern, and Hs_HSC_Toren. These stemness gene sets are presented in Supplementary Table S2.

### Single-sample gene-set enrichment analysis

We calculated the enrichment score of a gene set, which represents a stemness signature, biological process, pathway, or phenotypic feature, in a tumor bulk or single cancer cell by the single-sample gene-set enrichment analysis (ssGSEA) [40]. The ssGSEA calculates a gene set’s enrichment score in a sample based on their expression profiles. We implemented the ssGSEA algorithm with the “GSVA” R package. The gene sets analyzed are presented in Supplementary Table S2.

### Clustering analysis

We used hierarchical clustering to identify stemness subtypes of cancers based on their enrichment scores of the six stemness signatures. We performed hierarchical clustering analysis using the R package “hclust” with the parameters: method = “ward.D2” and members = NULL. Prior to clustering, the data of ssGSEA scores were transformed by z-score and translated into distance matrices by the “dist()” function with the parameter: method = “Euclidean.”

### Survival analysis

We utilized the Kaplan-Meier (K-M) model to compare overall survival (OS), disease-specific survival (DSS), disease-free interval (DFI), and progression-free interval (PFI) among different subgroups of cancer patients. Log-rank tests were utilized to assess the significance of their differences. We implemented the analyses with the function “survfit()” in the R package “survival.” Furthermore, we performed multivariate survival analysis by the Cox proportional hazards model to look into the correlations of the signatures of ESCs and HSCs, and the ratio of HSCs/ESCs with survival prognosis in pan-cancer after correcting for confounding variables, including age, sex, and tumor stage. The “age”, “ESCs”, “HSCs”, and “ratio of HSCs/ESCs” were continuous variables, and the “sex” (male versus female) and “tumor stage” (early (stage I-П) versus late (stage III-IV)) were binary variables. We performed the multivariate survival analysis with the function “coxph()” in the R package “survival.”

### Evaluation of genomic instability, ITH, and immune scores

Somatic mutations and copy number alterations (CNAs) reflect genomic instability. The tumor mutation burden (TMB) was defined as the total number of somatic mutations in the tumor. We obtained CNA scores in TCGA pan-cancer from the publication by Knijnenburg et al. [41]. We used GISTIC2 [42] to calculate G-scores in tumors with the input of “SNP6” files. The G-score reflects the amplitude of the somatic CNA and the frequency of its occurrence across a group of samples [42]. Besides, we utilized the DEPTH algorithm [19] to evaluate ITH in tumors, which measures ITH at mRNA level based on the heterogeneity of gene expression perturbations. In addition, we employed the ABSOLUTE algorithm [43] to evaluate tumor aneuploidy, namely ploidy scores, with the input of “SNP6” files. We calculated immune scores representing immune infiltration levels in tumors using the ESTIMATE algorithm [44] with the input of gene expression matrix.

### Identification of marker genes for subtypes

To identify marker genes for a stemness subtype, we first identified the upregulated genes in the subtype versus each of the other subtypes using the threshold of Student’s *t* test adjusted *P* value < 0.01 and mean expression fold change > 2. The marker genes for the stemness subtype were the common genes among the sets of upregulated genes.

### Identification of upregulated proteins for subtypes

To identify upregulated proteins in a stemness subtype, we compared the protein expression profiling in a stemness subtype with that in each of the other subtypes. The upregulated proteins for the stemness subtype were the common proteins identified as upregulated in the stemness subtype versus each of the other subtypes, with the threshold of Student’s *t* test adjusted *P* value < 0.05 and mean expression fold change > 1.

### Logistic regression analysis

We employed logistic regression models to predict tumors with higher immune scores (>median) versus those with lower immune scores (< median) with four predictors (ESC score, HSC score, CNA score, and TMB). In logistic regression analyses, all predictors’ values were normalized by z-score; the R function “glm” was utilized to fit the binary model with the parameter “family” as “binomial” and other parameters as default in “glm;” the standardized regression coefficients (β values) were obtained with the function “lm.beta” in the R package “QuantPsyc.”

### Statistical analysis

In class comparisons, for non-normally distributed data, we utilized Mann–Whitney *U* tests (for two classes) or Kruskal–Wallis (K–W) tests (for more than two classes); for normally distributed data, we used Student’s *t* tests (for two classes) or ANOVA tests (for more than two classes). In analysis of contingency tables, we utilized Fisher’s exact tests or Chi-square tests. In evaluating the correlations between the enrichment of molecular features and stemness scores, we employed the Spearman method. To adjust for *P* values in multiple tests, we utilized the Benjamini-Hochberg method [45] to calculate the false discovery rate (FDR). We implemented all statistical analyses in the R programming environment (version 4.2.2).

## Results

### Stemness signatures-based clustering analysis identifies four subtypes of pan-cancer

Based on the enrichment scores (ssGSEA scores) of the six stemness signatures, we identified four subtypes of pan-cancer by hierarchical clustering (Fig. 1A). We termed the four subtypes StC1, StC2, StC3, and StC4, respectively. StC1 displayed high enrichment of both ESC and HSC signatures. StC2 showed high enrichment of ESC but low enrichment of HSC signatures. In contrast to StC2, StC3 had low enrichment of ESC but high enrichment of HSC signatures. StC4 displayed the lowest enrichment of both ESC and HSC signatures. We defined the enrichment score of the ESC signature in a tumor as the average enrichment scores of the three ESC signatures in the tumor and the enrichment score of the HSC signature as the average enrichment scores of the three HSC signatures, termed ESC score and HSC score, respectively. As expected, ESC and HSC scores were significantly different among the four subtypes: StC1 > StC2 > StC3 > StC4 for ESC and StC3 > StC1 > StC2 > StC4 for HSC (K–W test, *P* = 0) (Fig. 1B). The ratios of HSC over ESC signatures, defined as HSC score/ESC score, were also significantly different among the four subtypes: StC3 > StC1 > StC2 > StC4 (*P* = 0) (Fig. 1B).

**Fig. 1.**
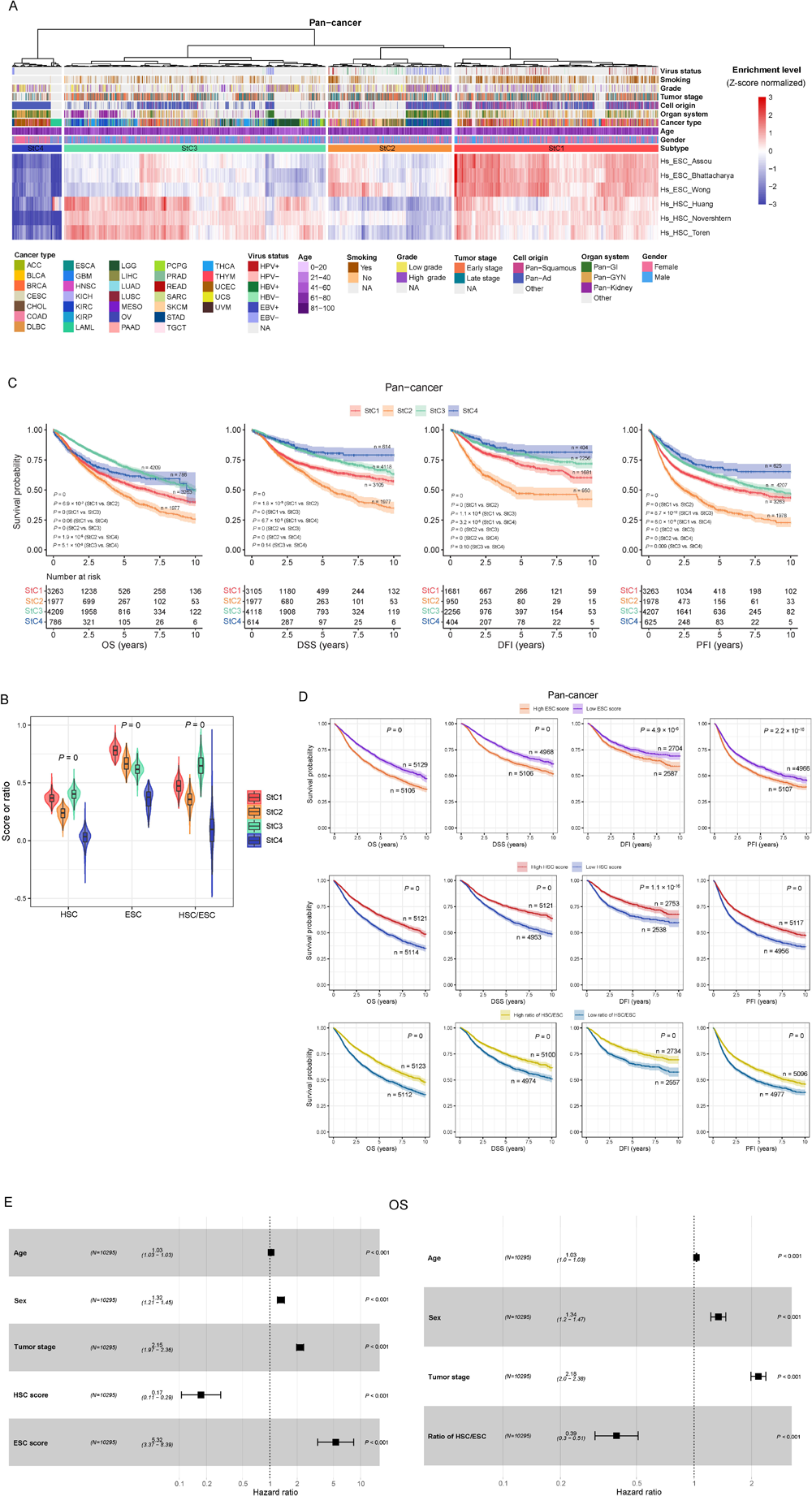
Identifying subtypes of pan-cancer based on stemness signatures. **A.** Hierarchical clustering identifies four stemness subtypes of TCGA pan-cancer: StC1, StC2, StC3 and StC4 based on the enrichment scores of six stemness gene sets, representing two types of stem cell backgrounds: embryonic stem cells (ESCs) and hematopoietic stem cells (HSCs). HPV: human papillomavirus, HBV: hepatitis B virus, EBV: Epstein–Barr virus, Ad: adenocarcinomas, GI: gastrointestinal, GYN: gynecological, NA: not available. **B.** Comparisons of HSC scores, ESC scores and ratios of HSC/ESC signatures among the four stemness subtypes of TCGA pan-cancer. The HSC score in a tumor is the average enrichment scores of the three HSC signatures. The ESC score in a tumor is the average enrichment scores of the three ESC signatures. The ratio of HSC/ESC signatures in a tumor is the ratio of HSC score over ESC score. **C.** Kaplan–Meier curves show that StC3 and StC2 likely have the best and worst 10-year OS, and that StC4 and StC2 likely have the best and worst 10-year DSS, DFI and PFI in pan-cancer. The log-rank test *P* values are shown. OS: overall survival, DSS: disease-specific survival, PFI: progression-free interval, DFI: disease-free interval. **D.** Kaplan–Meier curves show that the tumors with high ESC scores (> median) have significantly worse prognosis than the tumors with low ESC scores (< median) in all four 10-year endpoints (OS, DSS, DFI and PFI) in pan-cancer; In contrast, the tumors with high HSC scores or ratios of HSC/ESC signatures (> median) have significantly better prognosis than the tumors with low HSC scores or ratios of HSC/ESC signatures (< median) in all four 10-year endpoints in pan-cancer. The log-rank test *P* values are shown. **E.** Multivariate Cox proportional hazards regression analysis show that ESC scores have a significant inverse correlation with OS, and that HSC scores and the ratios of HSC/ESC signatures have a significant positive correlation with OS in pan-cancer. The “age”, “ESC score”, “HSC score”, and “ratio of HSC/ESC” are continuous variables, and the “sex” (male versus female) and “tumor stage” (early (stage I-П) versus late (stage III-IV)) are binary variables.

We compared four 10-year endpoints (OS, DSS, DFI, and PFI) among the four subtypes and found that they followed the pattern: StC3 > StC4 > StC1 > StC2 for OS and StC4 > StC3 > StC1 > StC2 for the other three endpoints (Fig. 1C). It indicates significant prognostic differences among these stemness subtypes. StC3 had the best OS; StC4 had the best PFI; StC2 showed the worst OS, DSS, DFI, and PFI; StC1 likely had the second worst OS, DSS, DFI, and PFI. Intriguingly, StC2 and StC3 exhibited an opposite pattern in expressing both types of stemness cell signatures, and the former had the worst prognosis, compared to the latter having the best OS prognosis. These results indicate that the expression of ESC signatures could be an adverse prognostic factor and the expression of HSC signatures be a positive prognostic factor. Indeed, we found that the tumors with high ESC scores (> median) displayed significantly worse prognosis than the tumors with low ESC scores (< median) in all four endpoints (*P* < 0.001) (Fig. 1D). In contrast, the tumors highly expressing HSC signatures showed significantly better prognosis than the tumors lowly expressing HSC signatures in all four endpoints (*P* < 0.001) (Fig. 1D).

Furthermore, the tumors with high ratios of HSC/ESC signatures had significantly better prognosis than the tumors with low ratios of HSC/ESC signatures in all four endpoints (*P* = 0) (Fig. 1D).

To explore whether the significant correlation between the enrichment of ESC and HSC signatures or ratio of HSC/ESC signatures and prognosis was confounded by other variables, such as age, sex, and tumor stage, we did multivariate (age, sex, tumor stage, and ESC and HSC scores or ratio of HSC/ESC signatures) survival analyses using the multivariate Cox proportional hazards model. This analysis showed that the ESC signature remained a significant risk factor and the HSC signature and ratio of HSC/ESC signatures protective factors in pan-cancer (Fig. 1E and Supplementary Fig. S1).

### Distribution of the stemness subtypes across individual cancer types

For individual cancer types, there were the highest proportions of BLCA, BRCA, CESC, COAD, HNSC, LUAD, LUSC, READ, TGCT, THYM, and UCS in StC1, the highest proportions of DLBC, ESCA, LIHC, OV, SKCM, STAD, and UVM in StC2, the highest proportions of ACC, CHOL, GBM, KICH, KIRC, KIRP, LGG, MESO, LUAD, PAAD, PCPG, PRAD, SARC, and THCA in StC3, and the highest proportions of LAML and UCEC in StC4 (Fig. 2A). Notably, most CESC (84.7%), LUSC (86.7%), TGCT (93.0%) and UCS (91.2%) tumors belonged to StC1, indicating the high enrichment of both ESC and HSC signatures in these cancer types.

**Fig. 2.**
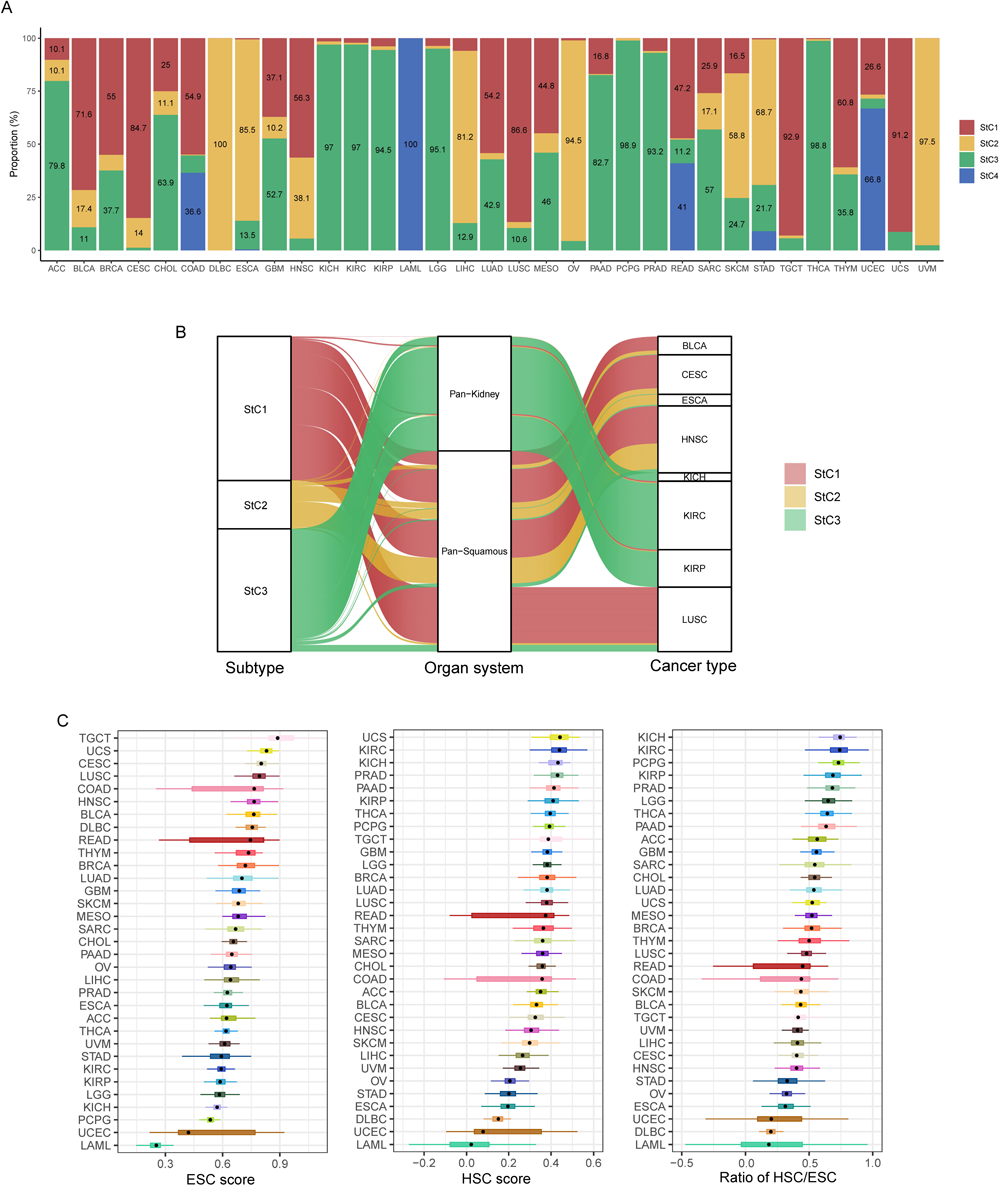
Distribution of the stemness subtypes across individual cancer types. **A.** Proportions of tumor samples belonging to each stemness subtype in each of the 33 TCGA cancer types. Only the proportions over 10% are shown in the bars. **B.** The Sankey diagram shows the stemness-subtype composition of pan-kidney and pan-squamous cell cancers. **C.** Comparisons of the enrichment scores of ESC and HSC signatures and their ratios across the 33 TCGA cancer types, ordered by the median value.

StC2 harbored all DLBC and most of ESCA (85.4%), LIHC (81.2%), OV (94.5%), and UVM (97.5%) tumors, suggesting the high enrichment of ESC but low enrichment of HSC signatures displayed in these cancer types. In contrast, StC3 harbored most of KICH (97.0%), KIRC (97.0%), KIRP (94.5%), LGG (95.1%), PAAD (82.7%), PCPG (98.9%), PRAD (93.2%), and THCA (98.8%) tumors, suggesting the low enrichment of ESC but high enrichment of HSC signatures shown in these cancer types. Finally, StC4 harbored all LAML cases, indicating the low enrichment of both ESC and HSC signatures in this cancer type. Interestingly, a majority (around 70%) of squamous cell carcinomas, such as CESC, HNSC, and LUSC, were classified into StC1 (Fig. 2B), suggesting the high stemness of squamous cell carcinomas. Another interesting finding was that most (96.2%) of kidney cancers, including KIRC, KICH, and KIRP, were grouped into StC3 (Fig. 2B). It suggests that kidney cancers have high HSC signatures but low ESC signatures.

We further compared the enrichment scores of ESC and HSC signatures and their ratios across the 33 TCGA cancer types (Fig. 2C). The analysis demonstrated that LAML, UCEC, PCPG, KICH, LGG, KIRP, and KIRC had the lowest median ESC scores, as compared to TGCT, UCS, CESC, LUSC, COAD, HNSC, and BLCA having the highest median ESC scores. Meanwhile, LAML, UCEC, DLBC, ESCA, STAD, and OV had the lowest median HSC scores, while UCS, KIRC, KICH, PRAD, PAAD, KIRP, and THCA had the highest median HSC scores. The ratios of HSC/ESC signatures were the lowest in LAML, DLBC, UCEC, ESCA, OV, and STAD and the highest in KICH, KIRC, PCPG, KIRP, PRAD, LGG, and THCA. This analysis showed that the cancer types with more excellent prognosis, such as kidney cancer, prostate cancer, thyroid cancer, and low-grade gliomas, had high enrichment of the HSC signature as well as ratio of HSC/ESC signatures. Contrastively, upper gastrointestinal cancer (such as esophageal, gastric, and liver cancer) and several gynecologic cancer (such as endometrial and ovarian cancer) displayed low enrichment of the HSC signature and ratio of HSC/ESC signatures.

### Correlates of the stemness subtypes with clinicopathologic features in pan-cancer

Among the four stemness subtypes, the proportion of early-stage (stage I-II) tumors followed the pattern: StC3 (74.0%) > StC4 (73.0%) > StC1 (52.4%) > StC2 (29.2%) (Chi-square test, *P* = 3.0 × 10^-59^) (Fig. 3A). Furthermore, the proportion of low-grade (G1-2) tumors followed the pattern: StC3 (50.21%) > StC4 (50.13%) > StC2 (48.3%) > StC1 (35.6%) (Chi-square test, *P* = 4.4 × 10^-13^) (Fig. 3A). These results overall conform to the prognostic difference among these subtypes. We further explored correlates of the stemness subtypes with therapeutic responses in pan-cancer. We observed that StC4 had the highest response (complete or partial response) rate (94.4%) to chemotherapy (*P* = 3.0 × 10^-16^) (Fig. 3A). It is justified since StC4 has the lowest stemness signatures to facilitate chemotherapy response [46].

**Fig. 3.**
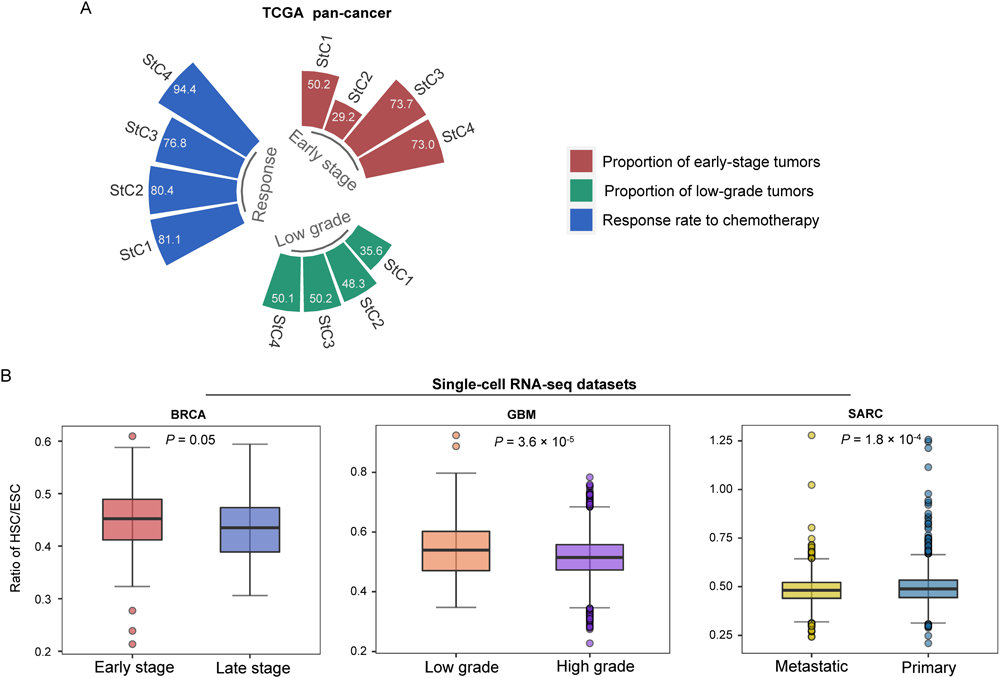
Comparisons of clinicopathologic features among the stemness subtypes of pan-cancer. **A.** The proportions of early-stage (stage I-II) tumors and low-grade (G1-2) tumors, and the response rates to chemotherapy among the four stemness subtypes in TCGA pan-cancer. Chi-square test, *P*L<L0.001. **B.** Comparisons of the ratios of HSC/ESC signatures between early-stage and late-stage cancer cells in BRCA, low-grade and high-grade cancer cells in GBM, and primary and metastatic cancer cells in SARC. The one-tailed Mann–Whitney *U* test *P* values are shown.

We further explored the association between stemness and clinicopathologic features in several cancer single-cell datasets. This analysis revealed that the ratios of HSC/ESC signatures were higher in cancer cells of early-stage tumors than in those of late-stage tumors in BRCA (*P* = 0.05) (Fig. 3B). In another cancer single-cell dataset for GBM, the ratios of HSC/ESC signatures were significantly higher in cancer cells of low-grade tumors than in those of high-grade tumors (*P* = 3.6 × 10^-5^) (Fig. 3B). Furthermore, the ratios of HSC/ESC signatures were significantly higher in cancer cells of primary tumors than in those of metastatic tumors in SARC (*P* = 1.8 × 10^-4^) (Fig. 3B). These results support the finding of the ratio of HSC/ESC signatures being a positive prognostic factor in tumors.

### Correlates of the stemness subtypes with molecular features in pan-cancer

Both increased TMB and CNAs reflects genomic instability [47]. We found TMB of the four stemness subtypes following the pattern: StC2 > StC1 > StC3 > StC4 (K–W test, *P* = 2.0 × 10^-279^) (Fig. 4A), and CNA scores following the pattern: StC2 > StC1 > StC4 > StC3 (K–W test, *P* = 6.3 × 10^-215^) (Fig. 4B). Furthermore, G-scores of copy number amplifications and deletions followed the same pattern as CNA scores: StC2 > StC1 > StC4 > StC3 (Fig. 4C). In addition, we found that ESC scores had significant positive correlations with both TMB and CNA scores in pan-cancer, and that HSC scores had significant negative correlations with both TMB and CNA scores (Fig. 4D). Furthermore, we found ESC and HSC scores to have a significant positive and negative correlation with microsatellite instability (MSI) scores evaluated by MSIsensor [48] in pan-cancer, respectively (*P* < 0.001) (Fig. 4D). These results collectively indicate that the enrichment of ESC and HSC signatures correlates positively and negatively with genomic instability, respectively. The ITH scores by DEPTH [19] were the highest in StC2 and the lowest in StC3 (*P* = 0) (Fig. 4E). It indicates that the enrichment of ESC and HSC signatures have a positive and negative correlation with ITH, respectively. Indeed, ESC scores displayed a significant positive correlation with ITH scores in pan-cancer (Spearman correlation, ρ = 0.13; *P* = 3.8 × 10^-33^), and HSC scores showed a significant negative correlation with ITH scores (ρ = -0.44; *P* = 0) (Fig. 4F).

**Fig. 4.**
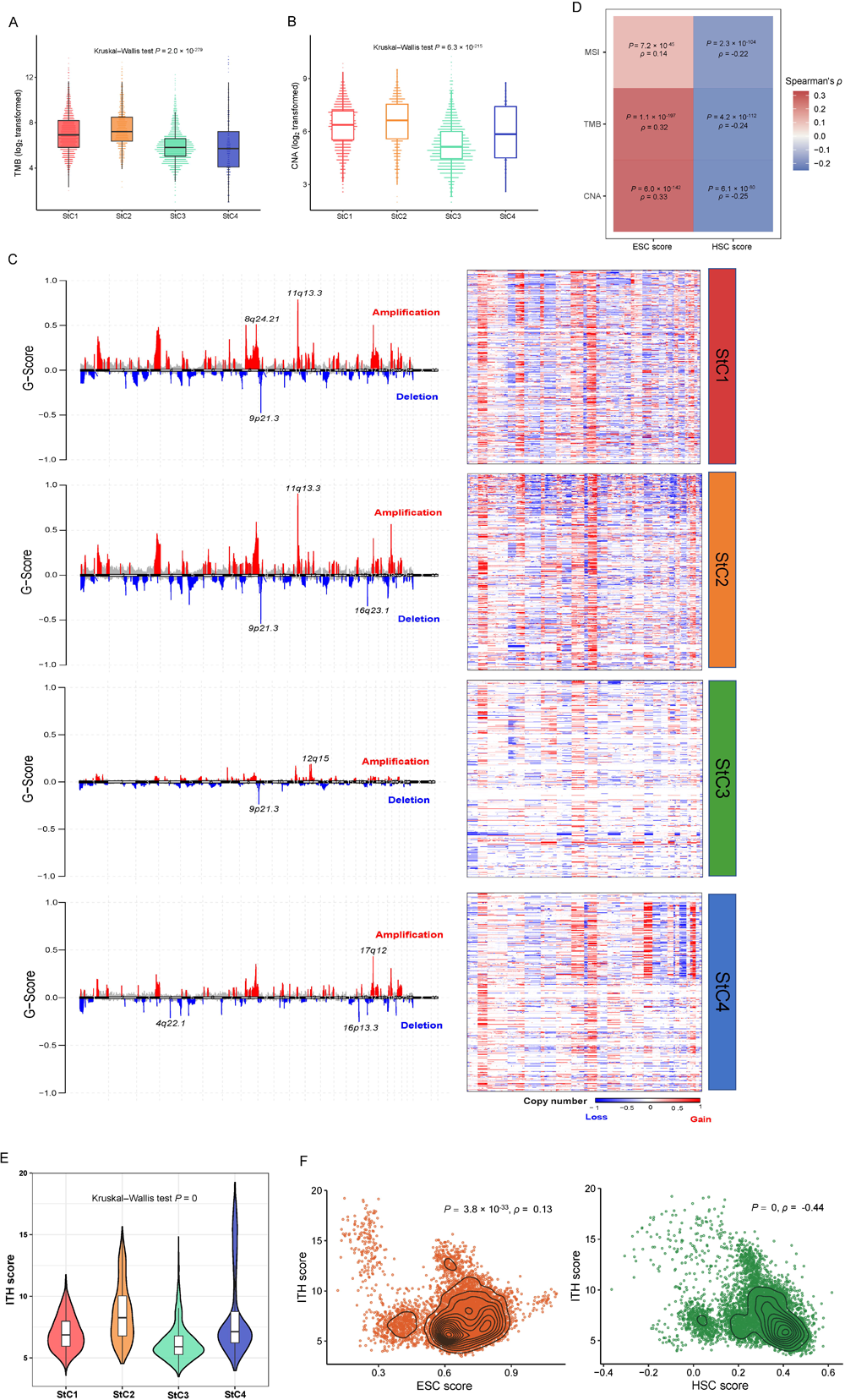
Comparisons of genome instability and intratumor heterogeneity among the stemness subtypes of pan-cancer. Comparisons of TMB (**A**), CNA scores (**B**), G-scores (**C**), and ITH scores (**E**) among the four stemness subtypes. The G-scores were calculated by GISTIC2 [42]. Heatmap (**D**) and scatter plots (**F**) show that ESC scores have significant positive correlations with MSI (**D**), TMB (**D**), CNA (**D**), and ITH scores (**F**), while HSC scores have significant negative correlations with MSI, TMB, CNA, and ITH scores in pan-cancer. TMB: tumor mutation burden, CNA: copy number alterations, ITH: intratumor heterogeneity, MSI: microsatellite instability. The Kruskal–Wallis test *P* values (**A, B, E**) and the Spearman correlation coefficients (ρ) and *P* values (**D, F**) are shown.

Furthermore, we compared the somatic mutation profiles among the four stemness subtypes. Fig. 5A displayed top ten genes having the highest mutation rates in each subtype. *TP53* showed the highest mutation rate (52%) in StC1 and the lowest mutation rate (20%) in StC3, although it was the most frequently mutated gene in StC3. *PTEN*, *PIK3CA*, *ARID1A*, *APC*, *KRAS*, and *FAT4* had the highest mutation rates in StC4 among the four subtypes.

**Fig. 5.**
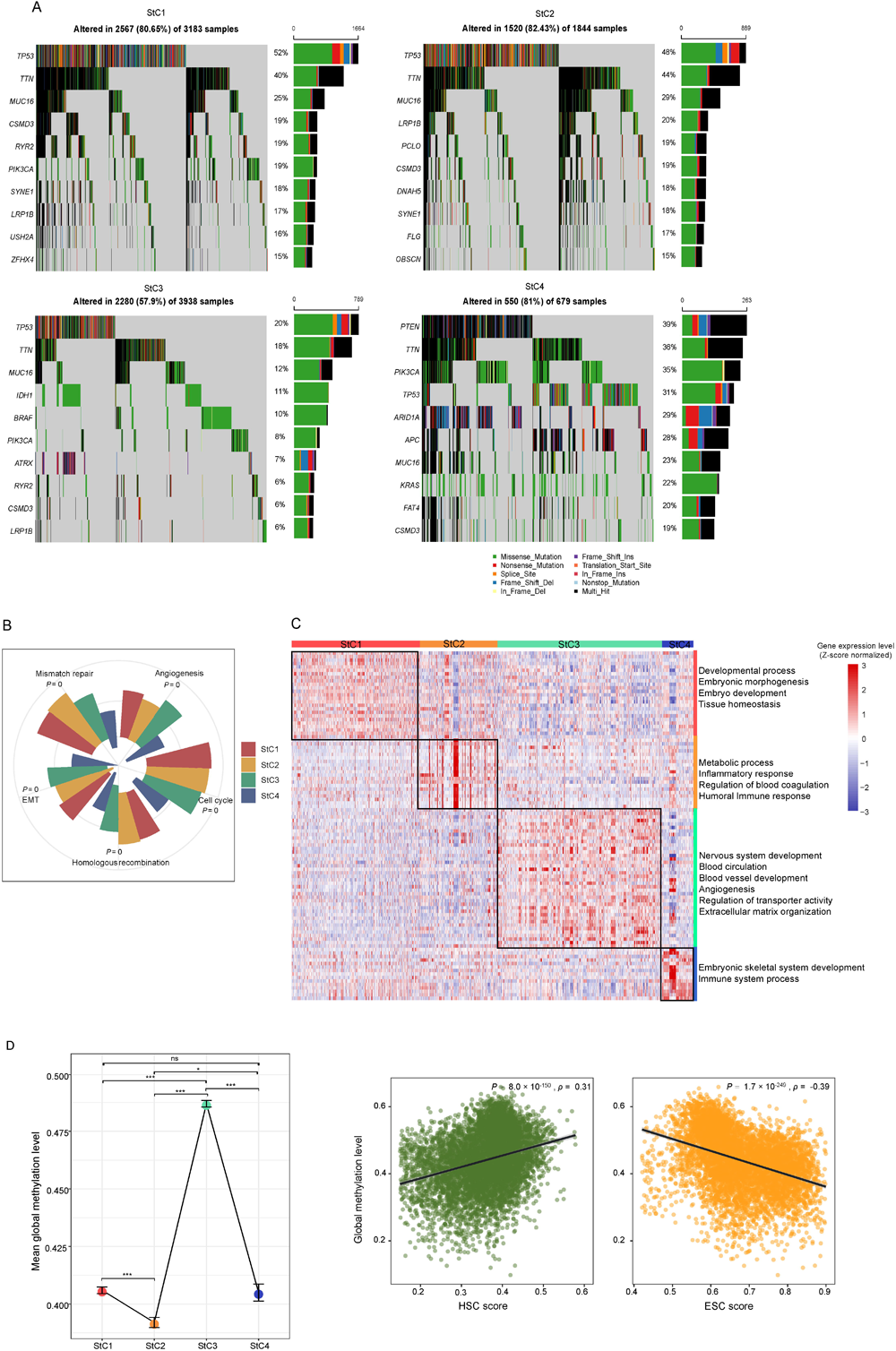
Comparisons of molecular features among the stemness subtypes of pan-cancer. **A.** Comparisons of somatic mutation profiles among the four stemness subtypes in pan-cancer. The oncoplot displays top ten genes having the highest mutation rates in each subtype. **B.** Comparisons of the enrichment scores of mismatch repair, homologous recombination, cell cycle, epithelial-mesenchymal transition (EMT) and angiogenesis signatures among the four stemness subtypes. The Kruskal–Wallis test *P* values are shown. **C.** Heatmap shows the expression levels of marker genes for each of the four stemness subtypes in pan-cancer (two-tailed Student’s *t* test, FDR < 0.01). The gene ontology (GO) biological process (BP) enriched in the stemness subtypes are shown on the right (FDR < 0.05). **D.** Left: comparisons of global methylation levels among the four stemness subtypes in pan-cancer. The mean global methylation level and error bar of each subtype are shown. Right: the scatter plots show that HSC scores have significant positive correlations with global methylation levels, while ESC scores have significant negative correlations with global methylation levels in pan-cancer. The one-tailed Mann–Whitney *U* test *P* values and the Spearman correlation coefficients (ρ) and *P* values are shown. * *P*L<L0.05, *** *P*L<L0.001, ^ns^ not significant.

Transcriptome analysis demonstrated that the enrichment scores of the mismatch repair, homologous recombination, and cell cycle signatures followed the pattern: StC1 > StC2 > StC3 > StC4 (*P* = 0) (Fig. 5B), consistent with the pattern of the enrichment scores of the ESC signature. It is justified as increased tumor stemness is linked to heightened genomic instability and cell proliferation potential in cancer [7].

In contrast, the enrichment scores of the epithelial-mesenchymal transition (EMT) and angiogenesis signatures followed the pattern: StC3 > StC1 > StC2 > StC4 (*P* = 0) (Fig. 5B), consistent with the pattern of the enrichment scores of the HSC signature. The positive association between the EMT and angiogenesis signatures and the HSC signature is imaginable since the HSC signature is enriched in the tumor stromal microenvironment. Transcriptome analysis also identified the marker genes for each of the four stemness subtypes (Fig. 5C and Supplementary Table S3). Based on the marker genes, we identified biological processes enriched in the subtypes using the GO database by g: Profiler [49]. StC1 was enriched for developmental process, embryonic morphogenesis, embryo development, and tissue homeostasis, in line with the high enrichment of both ESC and HSC signatures in this subtype. StC2 was enriched for metabolic process, inflammatory response, regulation of blood coagulation, and humoral immune response. StC3 was enriched for nervous system development, blood circulation, blood vessel development, angiogenesis, regulation of transporter activity, and extracellular matrix organization. StC4 was enriched for immune system process, and embryonic skeletal system development (Fig. 5C).

DNA methylation analysis revealed that global methylation levels [50] were the highest in StC3 and the lowest in StC2 (*P* < 0.001) (Fig. 5D). It suggests that the HSC and ESC signatures correlate positively and negatively with global methylation levels, respectively. We further confirmed this conclusion by Spearman rank correlation tests (Fig. 5D).

### Correlates of the ESC and HSC signatures with antitumor immune responses and immunotherapy responses

We found that HSC scores and ratios of HSC/ESC signatures correlated negatively with immune scores in pan-cancer (Spearman correlation test, *P* = 9.2 × 10^-124^ and 1.3 × 10^-119^, respectively) (Fig. 6A). Furthermore, the ratios of immunostimulatory to immunosuppressive signatures (pro-inflammatory/anti-inflammatory cytokines and M1/M2 macrophages) showed significant negative correlations with HSC scores and ratios of HSC/ESC signatures, while they had significant positive correlations with ESC scores (Fig. 6A). These results suggest that heightened enrichment of HSC and ESC signatures are associated with decreased and increased antitumor immune responses, respectively. Previous results have shown that the enrichment of HSC and ESC signatures has significant correlations with aneuploidy and TMB, which have been revealed to correlate significantly with antitumor immune responses [16]. Hence, the significant correlations between HSC and ESC signatures and antitumor immune responses could be associated their associations with aneuploidy and TMB. To explore this conjecture, we respectively predicted immune scores, ratios of pro-inflammatory/anti-inflammatory cytokines and ratios of M1/M2 macrophages (high (> median) versus low (< median)) using four variables: ESC score, HSC score, CNA score, and TMB, by logistic regression analyses. This analysis showed that ESC score and HSC score were a significant positive and negative predictor of ratios of pro-inflammatory/anti-inflammatory cytokines and ratios of M1/M2 macrophages, respectively; HSC score was a significant negative predictor of immune scores (βL=L-1.05; *P*L=L1.4L×L10^−46^) and ESC score a positive predictor of immune scores—albeit not significant (βL=L0.13; *P*L=L0.06) (Fig. 6B). These results suggest that the correlates of HSC and ESC signatures with antitumor immune responses are likely independent of their associations with aneuploidy and TMB.

**Fig. 6.**
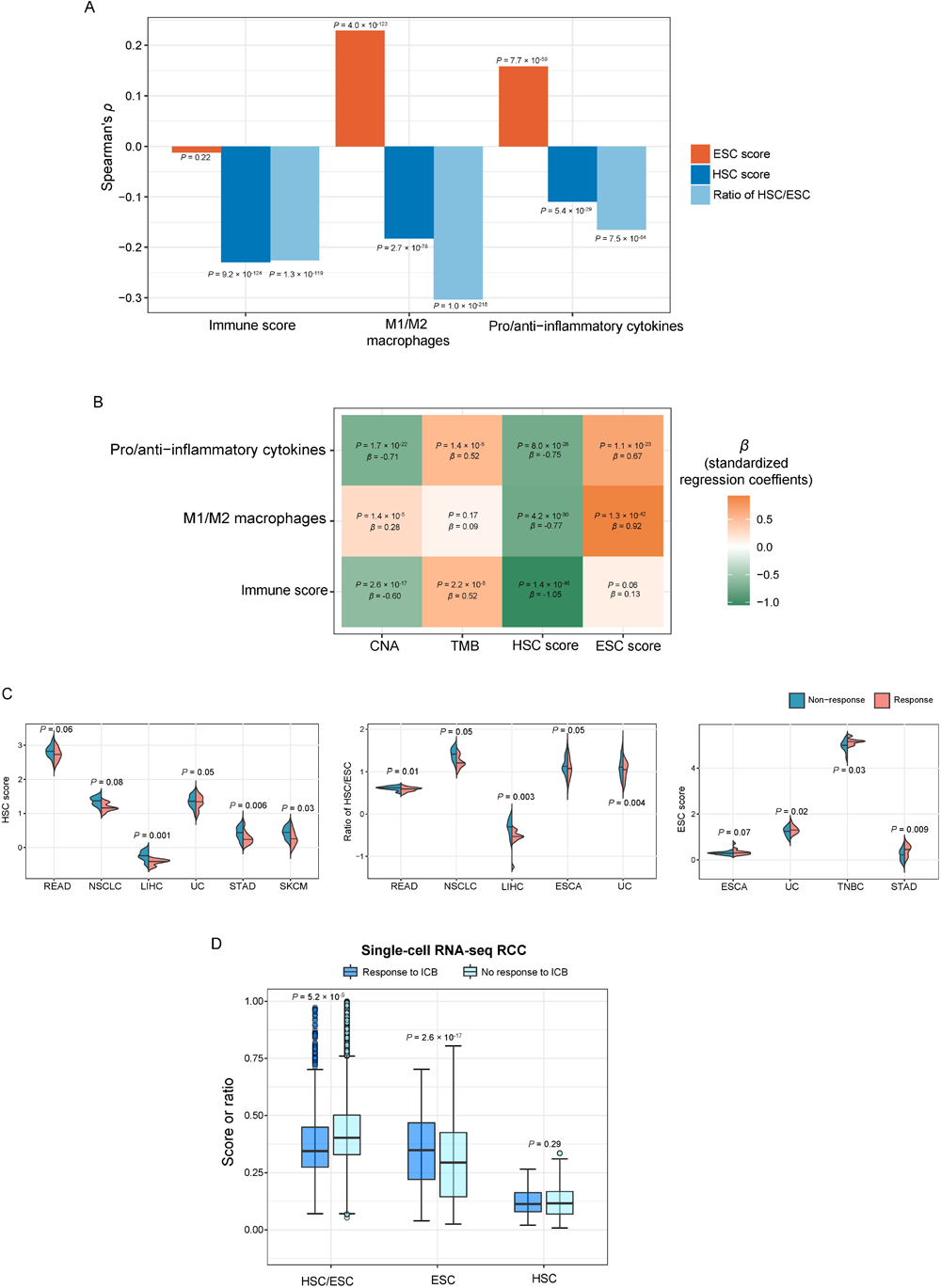
Correlations of ESC and HSC signatures with antitumor immune responses and immunotherapy responses. **A.** The bar charts show the correlations of ESC scores, HSC scores and ratios of HSC/ESC signatures with immune scores, M1/M2 macrophages and pro-inflammatory/anti-inflammatory cytokines in pan-cancer. The Spearman correlation *P* values are shown. **B.** Logistic regression analysis show that ESC score and HSC score are a positive and negative predictor of ratios of pro-inflammatory/anti-inflammatory cytokines, ratios of M1/M2 macrophages, and immune scores, respectively, after correcting for TMB and CNA. The standardized regression coefficients (β) and *P* values are shown. **C.** The correlations of HSC and ESC signatures and their ratios with anti-PD-1/PD-L1 immunotherapy responses in several cancer cohorts. NSCLC: non-small cell lung carcinoma, UC: urothelial cancer, TNBC: triple-negative breast cancer. The one-tailed Mann–Whitney *U* test *P* values are shown. **D.** The correlations of HSC and ESC signatures and their ratios with ICB responses in a scRNA-seq dataset for RCC. ICB: immune checkpoint blockade. The one-tailed Mann–Whitney *U* test *P* values are shown.

We further analyzed correlations of HSC and ESC signatures with anti-PD-1/PD-L1 immunotherapy responses in several cancer cohorts. We observed HSC scores to be likely higher in non-responsive than in responsive cancers in six cancer cohorts (*P* < 0.1) (Fig. 6C). Likewise, ratios of HSC/ESC signatures were likely higher in non-responsive than in responsive cancers in five cancer cohorts (*P* < 0.1) (Fig. 7C). However, ESC scores were likely higher in responsive than in non-responsive cancers in four cancer cohorts (*P* < 0.1) (Fig. 6C). In a single-cell transcriptome dataset for RCC [30], we also found ESC scores to be significantly higher in cancer cells of ICB-responsive tumors than in those of ICB-non-responsive tumors (*P* < 0.001) (Fig. 7D). In contrast, ratios of HSC/ESC signatures were significantly lower in cancer cells of ICB-non-responsive tumors than in those of ICB-responsive tumors (*P* < 0.001) (Fig. 7D), but there was no significant difference of HSC scores between ICB-responsive and ICB-non-responsive tumors (*P* = 0.29) (Fig. 7D). These results collectively suggest that the enrichment of HSC and ESC signatures are negatively and positively associated with immunotherapy responses, respectively.

**Fig. 7.**
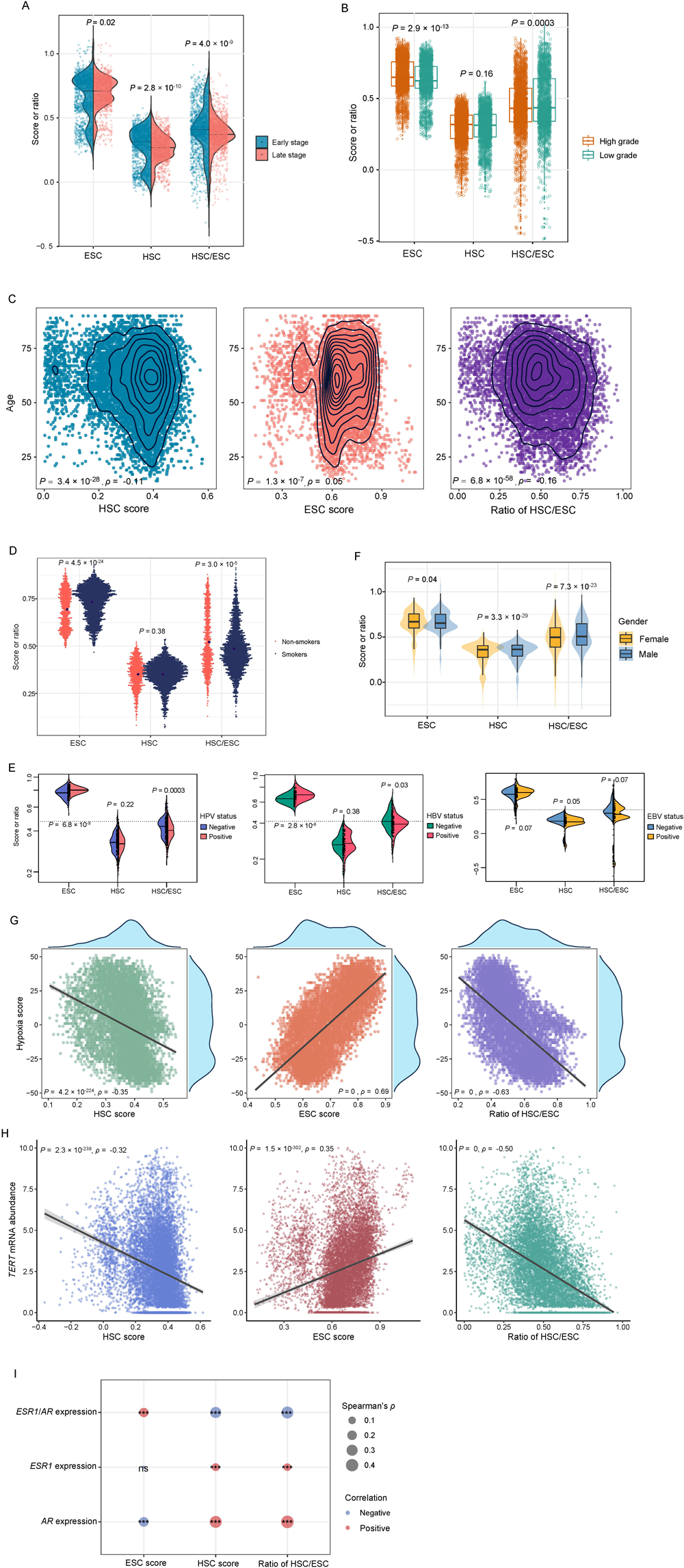
Correlations of ESC and HSC signatures with clinicopathologic and etiological factors in pan-cancer. Comparisons of HSC scores, ESC scores and ratios of HSC/ESC signatures between early-stage and late-stage (**A**), low-grade and high-grade (**B**), smoker and non-smoker (**D**), HPV/HBV/EBV-positive and -negative (**E**), and female and male (**F**) cancers, respectively. Correlations of HSC and ESC signatures and their ratios with age (**C**), hypoxia scores (**G**), *TERT* mRNA abundance (**H**), and the expression levels of *ESR1*, *AR* and their ratios (**I**). *ESR1*: estrogen receptor 1, *AR*: androgen receptor. The one-tailed Mann–Whitney U test *P* values are shown in (**A, B, D, E, F**), and the Spearman correlation coefficients (ρ) and *P* values are shown in (**C, G, H**). *** *P*L<L0.001, ^ns^ not significant.

### Correlates of the ESC and HSC signatures with clinicopathologic and etiological factors in pan-cancer

We found ESC scores to be significantly higher in late-stage than in early-stage cancers (*P* = 0.02) (Fig. 7A). In contrast, HSC scores were significantly lower in late-stage than in early-stage cancers (*P* = 2.8 × 10^-10^). The ratios of HSC/ESC signatures were also significantly lower in late-stage than in early-stage cancers (*P* = 4.0 × 10^-9^). In addition, ESC scores were significantly higher in high-grade than in low-grade cancers (*P* = 2.9 × 10^-13^), and HSC scores were likely lower in high-grade than in low-grade cancers (*P* = 0.16) (Fig. 7B). The ratios of HSC/ESC signatures were significantly lower in high-grade than in low-grade cancers (*P* = 2.9 × 10^-4^). Again, these results support that the ESC signature, HSC signature, and ratio of HSC/ESC signatures are a negative, positive, and positive prognostic factor in cancer, respectively.

Aging, tobacco, and viral infection [51–53] are common risk factors for cancer. Interestingly, ESC and HSC scores had a significant positive and negative correlation with ages in pan-cancer, respectively (Spearman correlation test, *P* < 0.001) (Fig. 7C). The ratios of HSC/ESC signatures showed a significant negative correlation with ages in pan-cancer (*P* = 6.8 × 10^-58^). Smokers displayed significantly higher ESC scores than non-smokers in pan-cancer (*P* = 4.5 × 10^-24^), while there was no significant difference in HSC scores between smoker and non-smoker cancers (*P* = 0.38) (Fig. 7D). The ratios of HSC/ESC signatures were significantly higher in non-smoker than in smoker cancers (*P* = 3.0 × 10^-5^). Viral infection is an important etiological factor for several cancers, such as human papillomavirus (HPV) for HNSC and CESC, hepatitis B virus (HBV) for LIHC, and Epstein–Barr virus (EBV) for lymphoma and nasopharyngeal cancers. We found that HPV/HBV/EBV-positive cancers had significantly higher ESC scores than negative cancers in pan-cancer, while the former group showed significantly lower HSC scores and ratios of HSC/ESC signatures than the latter group (*P* < 0.1) (Fig. 7E). In addition, females displayed significantly higher ESC scores but significantly lower HSC scores and ratios of HSC/ESC signatures than males in pan-cancer (*P* = 0.01, 1.3 × 10^-6^ and 1.6 × 10^-14^, respectively) (Fig. 7F).

Hypoxia is one of the fundamental properties of cancer and plays a key role in cancer development [54]. Of note, hypoxia scores showed a significant positive, negative, and negative correlation with ESC scores, HSC scores, and ratios of HSC/ESC signatures, respectively (Fig. 7G). *TERT* (telomerase reverse transcriptase) plays an important role in maintaining telomere structures and length [55]. We found *TERT* mRNA abundance having a significant positive, negative, and negative correlation with ESC scores, HSC scores, and ratios of HSC/ESC signatures, respectively (*P* < 0.001; ρ = 0.35, -0.32 and -0.50, respectively) (Fig. 7H). It indicates that the ESC-like signatures are associated with telomere lengthening in cancer cells while the HSC-like signatures are associated with telomere shortening. In addition, the expression of *ESR1* (estrogen receptor 1) and *AR* (androgen receptor) showed significant positive correlations with HSC scores and ratios of HSC/ESC signatures, while *AR* expression levels were negatively correlated with ESC scores (Fig. 7I). Furthermore, the ratios of *ESR1*/*AR* displayed a significant positive correlation with ESC scores and significant negative correlations with HSC scores and ratios of HSC/ESC signatures. This result is concordant with the previous finding of females having higher ESC scores and lower HSC scores and ratios of HSC/ESC signatures than males in pan-cancer.

### Validation of the stemness signatures-based subtyping method in proteomics data

Furthermore, we performed a hierarchical clustering analysis of pan-cancer based on the enrichment scores of the six stemness signatures in proteomics data [39]. Consistent with the result from transcriptomics data, we identified four subtypes of pan-cancer, also termed StC1, StC2, StC3, and StC4, respectively (Fig. 8A). Likewise, we identified the marker proteins for each of the four stemness subtypes (Supplementary Table S4). We also identified biological processes enriched in the subtypes based on the marker proteins using the R package “clusterProfiler” [56] (Fig. 8B). We found that StC1 was enriched for the biological processes representing active molecular activities and DNA repair, such as DNA replication, regulation of DNA repair, positive regulation of response to DNA damage stimulus, positive regulation of DNA metabolic process, RNA splicing, RNA catabolic process, RNA transport, regulation of translation, protein localization to nucleus, protein folding, and regulation of intracellular protein transport. It is in line with StC1 having high enrichment of both ESC and HSC signatures. StC2 was enriched for immune activities, including positive regulation of innate immune response, lymphocyte proliferation, antigen processing and presentation, and response to type I interferon. It is consistent with the result from transcriptomics analysis that StC2 is enriched for immune signatures. StC3 was enriched for regulation of angiogenesis, extracellular matrix organization, and regulation of wound healing, consistent with the result from transcriptomics analysis. StC4 was enriched for regulation of autophagy, positive regulation of insulin receptor signaling pathway, and endosomal transport, similar to the result from transcriptomics data.

**Fig. 8.**
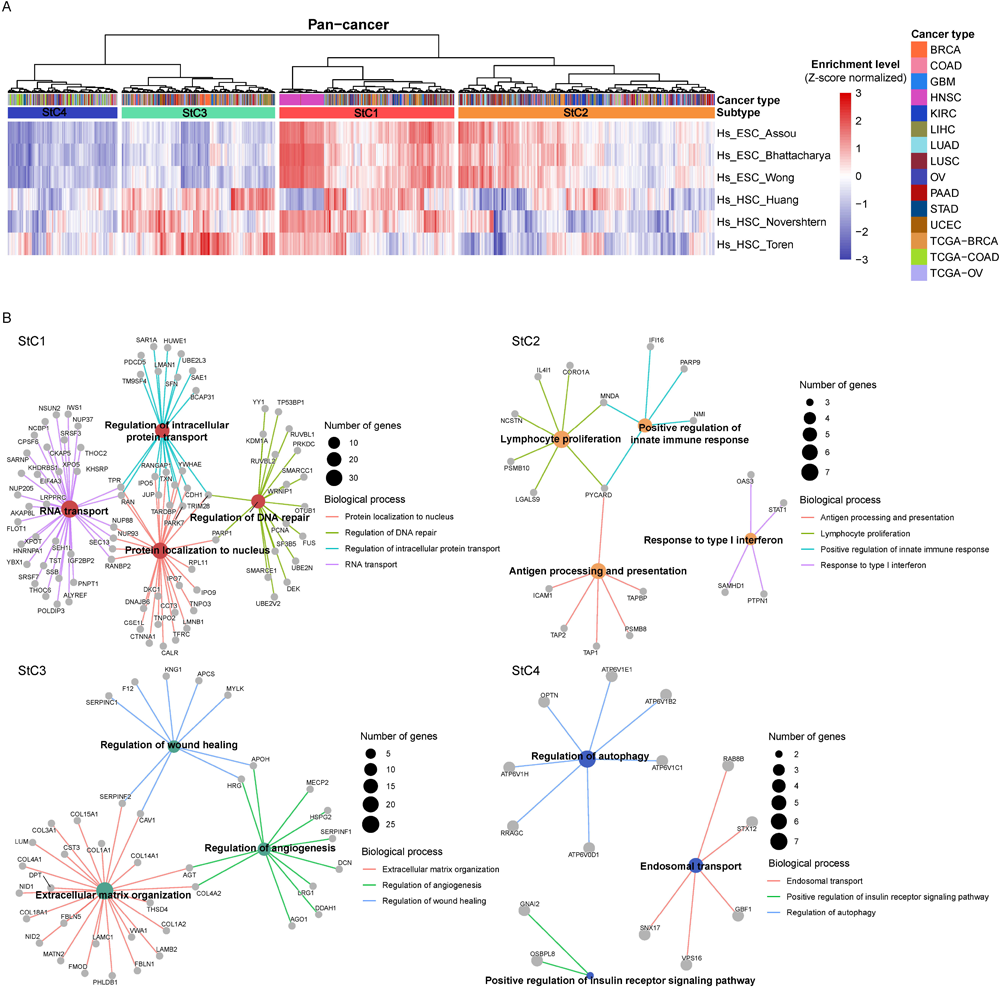
Validation of the stemness signatures-based subtyping method in proteomics data. **A.** Hierarchical clustering identifies four stemness subtypes (StC1, StC2, StC3 and StC4) based on protein expression profiles in pan-cancer. **B.** Enrichment plots show the biological processes enriched in the four subtypes using GO database by the R package “clusterProfiler.” Each protein-coding gene is shown as gray pie. The biological processes were identified using a threshold of FDR < 0.05.

Taken together, this analysis suggests that the stemness signatures-based subtyping method for pan-cancer is reproducible at the protein level.

## Discussion

This study identified four subtypes of TCGA pan-cancer based on the enrichment of six stemness signatures associated with ESCs and HSCs. The four subtypes exhibited significantly different molecular and clinical features. The subtype StC3, characterized by low enrichment of ESC but high enrichment of HSC signatures, has the best OS, the lowest aneuploidy level, ITH and *TP53* mutation rate, the highest enrichment of the EMT and angiogenesis signatures, and the highest global methylation levels. This subtype contains a majority of kidney, pancreatic, prostate, thyroid, and lower grade glioma tumors. In contrast, the subtype StC2, characterized by high enrichment of ESC but low enrichment of HSC signatures, has the worst prognosis, the highest aneuploidy level, tumor mutation loads and ITH, and the lowest global methylation levels. This subtype involves all DLBC and most of esophageal, liver, ovarian, and uveal melanoma tumors. StC1, characterized by high enrichment of both ESCs and HSCs signatures, has the second worst prognosis, the highest *TP53* mutation rate (52%), the highest enrichment of DNA mismatch repair, homologous recombination, and cell cycle signatures, and the second highest levels of genomic instability. StC1 harbors most squamous cell cancers, testicular germ cell tumors, and uterine carcinosarcomas. StC4 is characterized by the lowest enrichment of both ESC and HSC signatures; this subtype has the best progression-free survival, the highest response rate to chemotherapy, the lowest tumor mutation loads, the most frequent mutations in *PTEN*, *PIK3CA*, *ARID1A*, *APC*, *KRAS*, and *FAT4*, and the lowest enrichment of DNA mismatch repair, homologous recombination, cell cycle, EMT, and angiogenesis signatures. StC4 contains all acute myeloid leukemia cases.

This analysis further demonstrated that the ESC signature is a risk factor and that the HSC signature and ratio of HSC/ESC signatures are protective factors in cancer. Aging, smoking and viral infection likely confer higher enrichment of ESC signatures but lower enrichment of HSC signatures and ratio of HSC/ESC signatures on cancer.

A significant distinction between ESCs and HSCs is their different properties of cell divisions. ESCs expand themselves by symmetrical cell divisions [57], while HSCs maintain a balance between self-renewal and differentiation by asymmetric cell divisions [58]. Disruption of asymmetric cell divisions of stem cells may facilitate neoplastic transformation of stem cells [59]. Hence, the different prognostic correlates between the ESC and HSC signatures could be attributed to their distinct properties of cell divisions. It has been reported that several proteins play key roles in maintaining asymmetric cell divisions, including Par3, Par6, aPKC, NUMA, LGN, Gαi, and Prospero [60]. Of note, we found that the expression levels of the genes encoding these proteins likely had significant negative correlations with ESC scores but significant positive correlations with HSC scores and ratios of HSC/ESC signatures in TCGA pan-cancer. In addition, p53 dysfunction can promote a shift from asymmetric to symmetric cell divisions [61]. We observed that *TP53*-mutated tumors had significantly higher ESC scores but significantly lower HSC scores and ratios of HSC/ESC signatures compared to *TP53*-wildtype tumors in TCGA pan-cancer. These data support that the ESC and HSC signatures are associated with increased and reduced symmetric cell divisions in cancer. Indeed, *MKI67*, a marker for proliferation, shows a significant positive expression correlation with ESC scores and negative expression correlations with HSC scores and ratios of HSC/ESC signatures in TCGA pan-cancer (ρ = 0.50, -0.39, and -0.66, respectively).

An interesting finding is that HSC scores are negatively correlated with immune infiltration levels in cancer, while ESC scores are positively correlated with them. This finding appears inconsistent with reports from previous studies [7, 17, 23]. However, a previous report [14] demonstrates that solid tumors accumulate immunosuppressive hematopoietic lineages at the tumor sites. Another study suggests that activation of HSC signatures can promote immunosuppression within the pre-metastatic niche of tumors [62]. These data support to a certain degree our finding that high HSC scores or ratios of HSC/ESC signatures are associated with enhanced resistance to immunotherapy in tumors.

To conclude, this study performed a novel classification of pan-cancer in the context of HSC and ESC signatures and identified four stemness subtypes. Besides stemness features, these subtypes display different molecular and clinical characteristics, including genome integrity, ITH, methylation levels, tumor microenvironment, tumor progression phenotypes, chemotherapy and immunotherapy responses, and survival prognosis. The ESC signature is an adverse prognostic factor in cancer, while the HSC signature and ratio of HSC/ESC signatures are favorable prognostic factors. The HSC and ESC signatures-based subtyping of cancer provides new insights into tumor biology and has potential clinical implications for diagnosis, prognosis, and treatment of cancers.

## Data availability

The authors declare that all data supporting the findings of this study are available within the paper and its supplementary information files.

## Author contributions

**Jiali Lei**: Methodology, Software, Validation, Formal analysis, Investigation, Data curation, Visualization, Writing - original draft, Writing - review & editing. **Jiangti Luo**: Software, Formal analysis, Data curation, Visualization. **Qian Liu**: Software, Data curation. **Xiaosheng Wang**: Conceptualization, Methodology, Resources, Investigation, Writing - original draft, Writing - review & editing, Supervision, Project administration, Funding acquisition.

## Acknowledgments

Not applicable.

## Supplementary data

Table S1. A summary of the datasets analyzed.

Table S2. The gene sets representing stemness signatures and biological processes.

Table S3. Marker genes for each of the four stemness subtypes in TCGA pan-cancer.

Table S4. Highly expressed proteins in each of the four stemness subtypes of pan-cancer identified based on proteomics.

**Fig. S1. Correlations of ESC and HSC signatures with prognosis in pan-cancer.** Multivariate Cox proportional hazards regression analysis show that ESC scores have a significant inverse correlation with DSS, DFI and PFI, and HSC scores and the ratios of HSC/ESC signatures have a significant positive correlation with DSS, DFI and PFI in pan-cancer. The “age”, “ESC score”, “HSC score”, and “ratio of HSC/ESC” are continuous variables, and the “sex” (male versus female) and “tumor stage” (early versus late) are binary variables.

