## Supplementary figures and images for "Pan-cancer analysis reveals embryonic and hematopoietic stem cell signatures to distinguish different cancer subtypes"

### Supplementary Figure 1

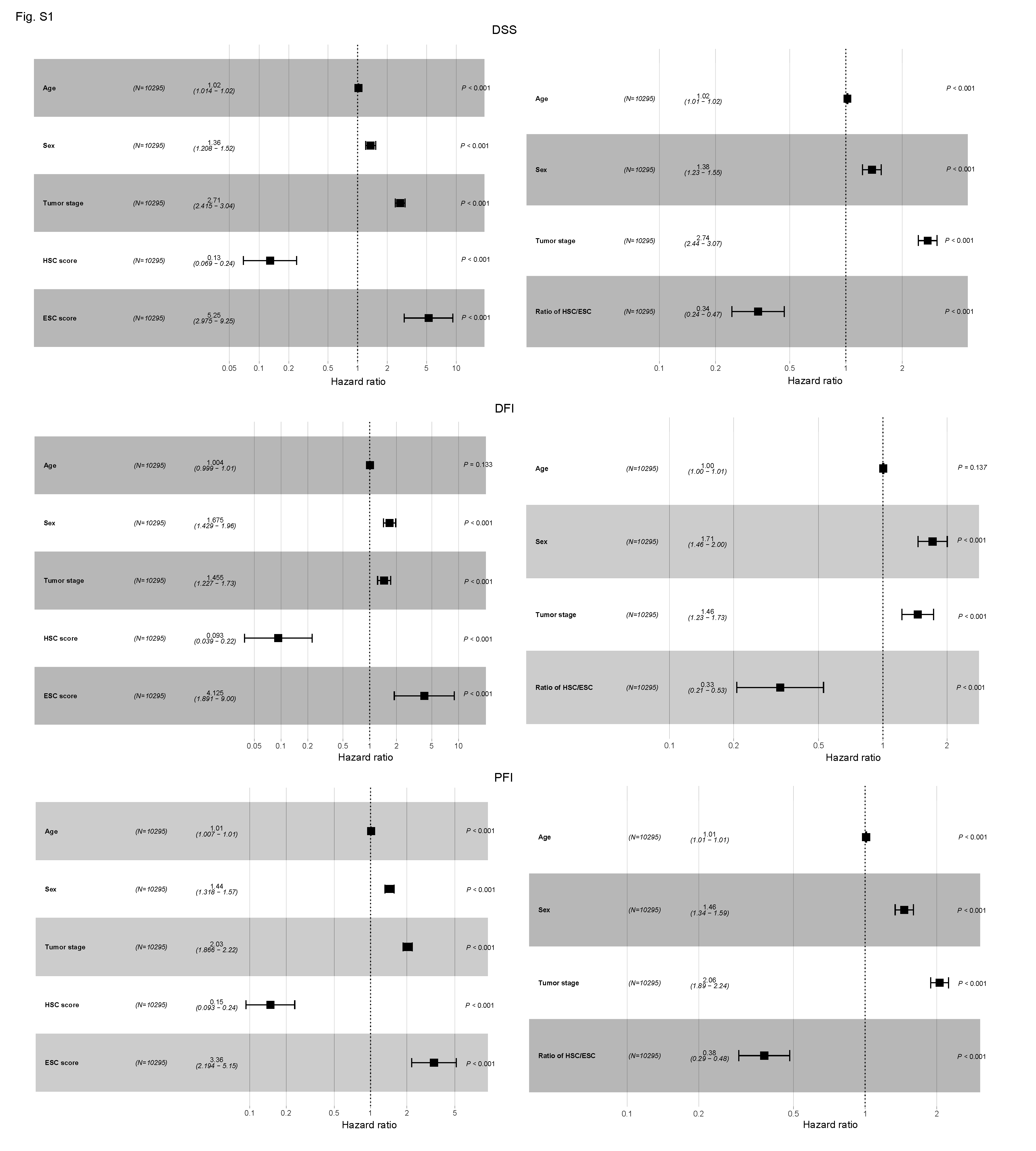
